# Automated and Simulation-Guided Multiplexed DNA-PAINT for Nanoparticle Characterization

**DOI:** 10.1101/2025.08.05.668442

**Authors:** Stijn van Veen, Emiel W. A. Visser, Lorenzo Albertazzi

## Abstract

Recent advances in lab automation have dramatically increased the throughput of material synthesis, enabling rapid screening of nanomaterials for specific applications in nanotechnology and nanomedicine. However, this progress highlights a key bottleneck: most high-resolution characterization techniques, such as electron microscopy, atomic force microscopy (AFM), and super-resolution microscopy, remain low-throughput and labor-intensive. To keep pace, characterization must evolve toward greater automation and scalability.

Here, we present an integrated and automatable workflow for multicolor DNA point accumulation for imaging in nanoscale topography (DNA-PAINT) super-resolution microscopy tailored for nanoparticles. Our approach combines kinetic simulations, automated multiplexed imaging, and streamlined image analysis to enable end-to-end automation, from experimental design to quantitative data output.

Simulations predict optimal experimental conditions, thus reducing the need for manual optimization. A fluidics system paired with a TIRF microscope is used to automate multiplexed imaging by rounds of imaging and probe exchange (exchange-PAINT) on multiple memorized positions without human oversight during the acquisition process. Finally, an image analysis pipeline tailored for NPs allows for the quantification of nanoparticle size and multiplexed ligand functionalization.

This methodology improves the throughput and reproducibility of single-molecule localization microscopy (SMLM) using DNA-PAINT and lowers the entry barrier for non-expert users, thus paving the way for broader adoption in nanomedicine and materials discovery workflows.

## Introduction

Materials science is undergoing a rapid evolution through an increase in automation, with the promise of self-driving laboratories.^1,2^ In this context, specialized hardware, such as pipetting robots or fluidic devices, has enabled vastly increased throughput in formulation, while potentially incorporating bulk analysis and recursive feedback to automatically generate new formulations. This increased throughput in formulation poses an immediate challenge for analytical techniques, as these materials need to be characterized and tested to identify the best-performing formulations. While simple analytical techniques are amenable to this, more advanced methods such as mass spectrometry, electron microscopy, and super-resolution optical microscopy are generally complex and cumbersome, and only a limited number of samples can be measured. Bringing advanced imaging techniques to automated pipelines can generate more information about the materials formulated and, therefore, support materials discovery. Consequently, there is a general interest in increasing the automation and throughput in advanced imaging.

Among the advanced imaging techniques used for materials characterization, super-resolution fluorescence microscopy is gaining momentum as it offers sub-diffraction imaging resolution,^3–7^ multicolor imaging, and single-molecule and single-particle sensitivity. One such technique, point accumulation for imaging in nanoscale topography (PAINT), has proven particularly promising for materials imaging thanks to its high-resolution, down to a few nanometers, and quantitative nature.^8–10^ PAINT uses fast and transiently binding fluorophore conjugates as an alternative to stochastic photo-activation to enable simultaneous imaging of multiple fluorescent events.^8^ Here we employ a specific variant, DNA-PAINT, which relies on transient interactions between two complementary single-stranded DNA oligonucleotides (ssDNA).

A key strength of DNA-PAINT lies in its simplicity and adaptability. It does not require complex buffer systems.^11^ or specialized equipment to induce photoswitching^12^, making it relatively easy to implement. Moreover, due to the transient binding, the imager strands can easily be flowed in and out, opening the way for multiplexed imaging that is not limited by the number of spectrally distinct dyes available.^8,13^ This makes DNA-PAINT an attractive option for multi-target imaging and characterization. Lastly, the binding kinetics for any given combination of complementary strands are predictable and can be quantified. This can be used to calculate the number of docking sites from the obtained timetraces, the imager concentration, and the experimentally determined kinetic constants.^10,14,15^ This has been used to quantify the amount of DNA on a nanoparticle,^15^ or even the orientation of antibodies on nanoparticles^16^.

These advantages come at a cost, however, such as the time-intensive nature of DNA-PAINT. Because super-resolution images rely on collecting data from thousands of transient binding events, imaging requires the acquisition of many frames, which becomes a bottleneck for high-throughput analysis. Moreover, the kinetic calculations are extremely sensitive to sample conditions, as they require both good statistics and for all events on a single particle to be temporally separated. This necessitates careful optimization of imaging conditions, especially for longer, multiplexed measurements, making the process labor-intensive. Lastly, while the transient nature allows for multiplexed imaging through fluid exchange, manual fluid handling slows down acquisition and complicates returning to the same field of view.

Here we present a streamlined workflow designed to significantly increase the throughput of DNA-PAINT, aligning it with recent advances in high-throughput sample preparation and analysis.

We leverage the predictability of DNA-PAINT’s binding kinetics, which follow Poisson statistics,^17^ to simulate synthetic timetraces and generate randomized distributions of binding events to optimize experimental parameters in advance, dramatically reducing the need for trial-and-error optimization. Through an automated flow system with microscope control, our approach enables continuous acquisition of kinetic data from nanoparticles without requiring user intervention once measurements begin.

This represents a shift from labor-intensive single-sample imaging to scalable, reproducible, and accessible single-molecule analysis. Ultimately, this approach is intended to increase the throughput of DNA-PAINT and establish it as a viable characterization method for a wider range of studies in the context of self-driving labs and machine-guided materials discovery.

## Results & Discussion

The goal of this research is to develop an automated workflow that can be applied to the characterization and quantification of materials using DNA-PAINT microscopy. After identifying a sample of interest and conjugating ssDNA to a target, the workflow has three key steps (Fig.1):

**Figure 1.**
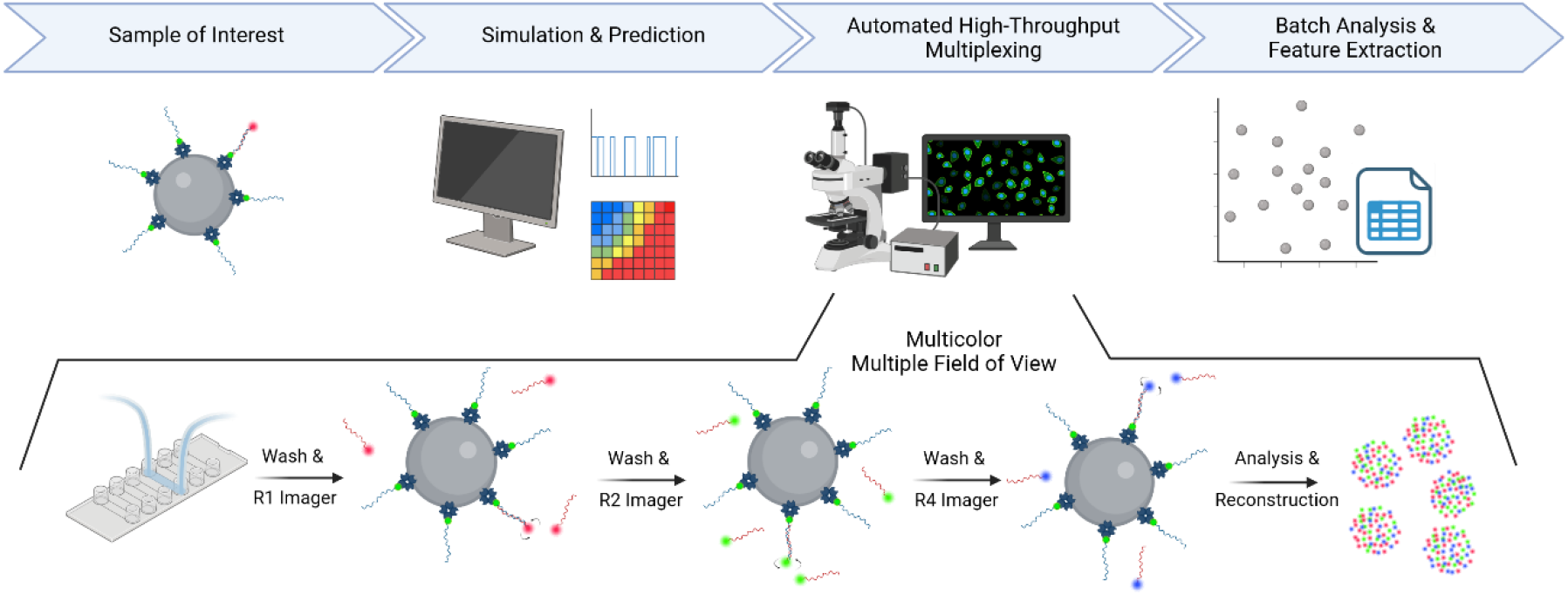
An overview figure displaying the sequential steps involved in the workflow. First, we identify our sample of interest and the likely degree of functionalization. Next, we employ timetrace simulations to optimize experimental parameters for high accuracy and precision. We automate the imaging process through the use of microfluidics. Lastly, all single-molecule data is aligned and analyzed to extract features like size and target count.

**Figure 2.**
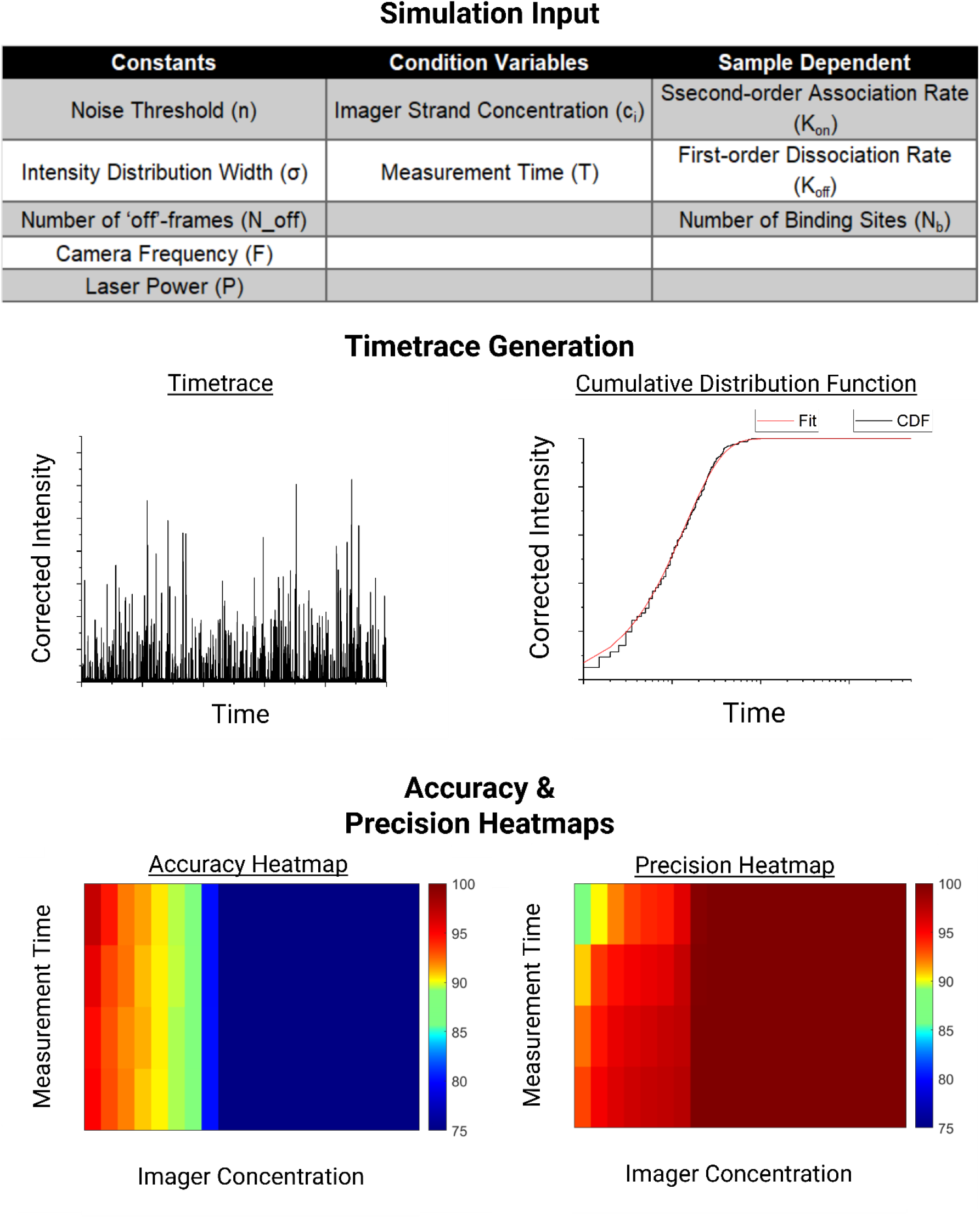
Overview of simulation input and output. The input variables are separated into constant variables that are the same for each simulation, experimental variables that depend on the sample (e.g., the DNA strand combination used), and the condition variables that are varied to find the optimal condition. An example timetrace and its corresponding cumulative distribution function fit are shown. The final output shown is an accuracy and precision heatmap based on 1000 simulated timetraces for many combinations of measurement time and imager concentration.

i. Simulating binding kinetics for different concentrations and timescales to determine optimal experimental conditions without requiring iterative trial-and-error;
ii. Executing fully automated multiplexed DNA-PAINT imaging using fluid exchange, for multiple fields of view;
iii. And processing image data using a batch analysis pipeline tailored to extract structural and functional nanoparticle information.

All protocols and code are freely available and designed for adaptability, aligning with the principles of open microscopy and broadening accessibility. Each of these steps is described in detail in the following sections.

To validate the robustness of the pipeline, we use commercially available streptavidin-coated polystyrene nanoparticles functionalized with biotin-ssDNA as a model system. This model system can be substituted with any DNA-functionalized platform, as shown in previous research that employed DNA conjugated to proteins targeting different antibody regions, for example,^16^ or the liposome imaging using cholesterol-DNA, presented later in this work.

Traditionally, it is challenging to quantify the degree of functionalization at the single-particle level. Techniques such as absorbance ratios or SPR provide ensemble-averaged data and do not allow for direct counting or give an accurate representation of the particle-to-particle heterogeneity within the population. In contrast, qPAINT enables direct molecular counting on a per-particle basis. This is a key distinction, as functionality can have a strong dependence on the degree of functionalization, which means averages are wholly insufficient in terms of information. It is very possible to have large differences within a population of particles, or even to have multiple different populations where only a specific subset shows the intended performance. Thus, molecular counting—when combined with the nanometer resolution and multiplexing capability of PAINT—offers a powerful and comprehensive approach to nanoparticle characterization.

### Simulation-Guided Optimization of Imaging Parameters

The molecular counting capability of DNA-PAINT is a powerful tool,^18,19^ but it is also very sensitive to sample conditions, particularly imager concentration. The qPAINT methodology relies on the predictable kinetic information, the binding and unbinding of ssDNA, to obtain the number of docking sites. Mathematically, this is represented by the following equation: *Nb* = *1/(k*_*on*_ *c*_*i*_ *τ*_*d*_), where Nb is the number of binding sites, k_on_ is the second-order association rate, c_i_ is the imager strand concentration, and τ_d_ is the true mean dark obtained experimentally from the cumulative distribution function of the individual dark times. Dark times refer to the time between binding events of the imager, where no fluorescent signal is present. The number of binding sites and the association rate depend on the sample used, while the imager concentration can be controlled and is a key factor in the density of events, which directly impacts the precision and accuracy of the qPAINT calculation.

To eliminate the need for tedious trial-and-error optimization, we aim to identify the optimal sample conditions beforehand. Since binding kinetics are predictable, they can be simulated in silico as shown by Jungmann et al.^8^ Whereas Picasso simulates image data, our approach focuses purely on generating the fluorescent timetraces and the subsequent qPAINT analysis, using custom MATLAB scripts with several input variables and constants, previously developed within the group.^20^ The step-by-step explanation of the timetrace simulation process is provided in the Materials and Methods.

Simulated binding timetraces demonstrate that small differences in concentration can lead to drastically different distributions and CDF fits. The strands used have relatively fast kinetics (Supporting Table 1), which is reflected in the high event density at an imager concentration of 1 nM, for example. This means there is a risk of double events, where two binding events happen at the same time point, leading to a risk of undercounting. At 0.5 nM, the fit distinctly improves, though the timetrace is still relatively dense, and double events could occur, albeit much less likely. If we further reduce the concentration to 0.1nM, we see that the timetrace becomes much more sparse, though the fit seems decent at first glance. However, low event counts reduce statistical robustness at this range.

To study this further, we run these simulations 1000 times for each combination of measurement time (T) and imager concentration (c_i_). Then we use these results to calculate the accuracy and precision. This gives us a better indication of how likely we are to retrieve the ground truth (Nb). We can see that for 200 R1-docking sites, 1nM is too high as suspected, and in general, the accuracy drops above 0.1 nM concentration. However, the lower statistics are reflected in the lowered precision at 0.1 nM concentration. Due to fewer binding events, there’s much more variance at shorter measurement times. Therefore, if we want to measure with both high accuracy and precision, while keeping our measurements short to guarantee high-throughput, then a concentration of 0.5 nM would be better than 0.1 nM. That is not something easily inferred from a single timetrace, nor easily identified by eye, underlining the nuance in deciding on the optimal parameters for an experiment.

For each DNA strand pair used in this article, accuracy and precision heatmaps were obtained for several different amounts of binding sites, which can be found in the Supporting Information (Supporting Figures 1-3). These were used to optimize the experiments displayed in the rest of this section.

### Automated Fluidic Multiplexing for Exchange-PAINT

As previously mentioned, DNA-PAINT is suitable for multiplexed imaging due to two main advantages:

i. The large library of orthogonal DNA sequences that can be used to tag structures of interest,^21,22^ and
ii. Its transient binding nature, which means that imager strands can be flowed out and replaced with a different imager without changing the sample, allowing sequential imaging.

Here we build on the exchange PAINT demonstrated by Jungmann et al,^23^ to design a high-throughput DNA-PAINT imaging platform.

To achieve this, we employ an automated platform incorporating both a microfluidic setup as well as an automatable microscope. Compared to manual fluid exchange, this process requires no user input after initial setup and allows for more thorough fluid displacement. The settings, instrumentation, and an example workflow are listed in the Materials and Methods, but the specific instrumentation used can be substituted, as long as a few criteria are met.

- First, precise time control is a necessity. A continuous multi-acquisition would not suffice, as the setup must allow for pauses between imaging steps, so that the flow steps and imaging steps are temporally separated.
- Secondly, a pump with multiple input lines is needed, whether that is through a valve system as used here or a design that allows several inlets into the same channel.
- Proper control over flow speed is important as well, to ensure that the sample remains intact after flow steps.
- Additionally, to achieve good throughput, automated position control is needed to image multiple fields of view sequentially.
- Lastly, fiducial markers are an absolute necessity to correct for drift, not only during the measurements, but also to account for drift in between imaging and flow steps to align the different markers in the final analysis.

All of the materials and flow parameters used are listed in the Materials and Methods.

To demonstrate the potential of this setup, we performed a 4-color DNA-PAINT experiment without human intervention. After selecting experimental conditions based on the simulations described in the previous section, we set up a measurement on NPs bearing three different DNA dockings. Over a span of 3.5 hours, 3 50×50 μm fields of view were imaged 4 times, once for each color, plus a repeat as quality control, each containing multiple particles. The R1 DNA pair was repeated in this measurement to verify that the flow steps do not affect the DNA functionalization of the particles (see quantification in Supporting Figure 4).

As shown in Figure 3, we observe that imagers can be flowed out and replaced as intended. This is evident from the localization count returning to the baseline after flow steps, indicating that the imager is fully removed and there is no cross-talk between channels. Additionally, the timetraces observed align well with the simulated timetraces at this concentration and DNA density, as shown in Supporting Figure 5.

**Figure 3.**
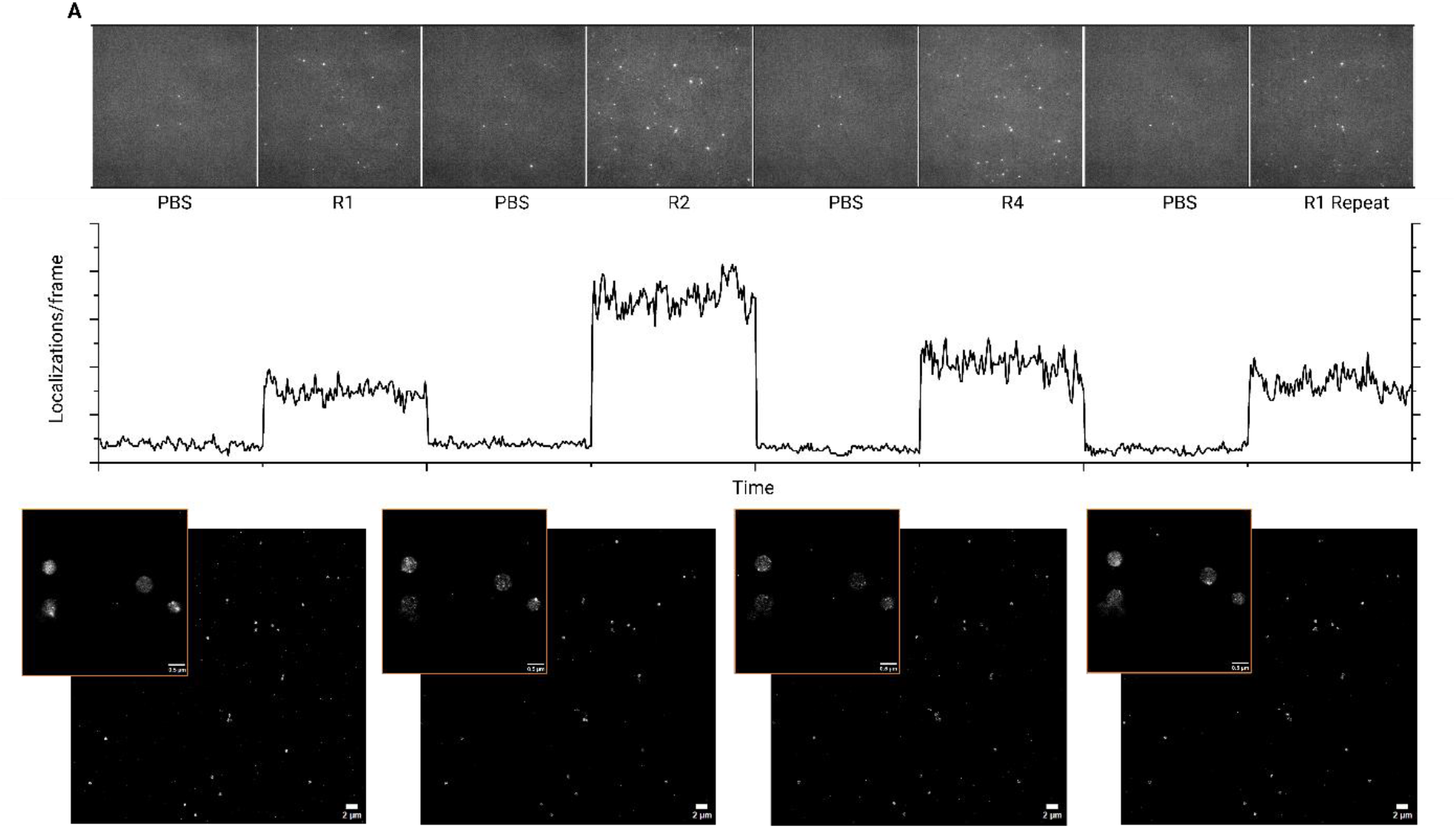
Visual representation of a multiplexed imaging experiment, with the first imager repeated at the end to show the consistency and reproducibility of the method. The top row displays a single representative frame from each step. The middle row displays the localization count during each step, which shows the effective removal of the imager strand in each PBS washing step. Important to note is that the sample includes some fiducials, which account for most of the remaining signal observed after washing. The bottom row shows the reconstruction for this field of view, with a cut-out showing representative particles. The data corresponding to this measurement is also displayed in (Supporting Figure 4).

### Automated Analysis and Feature Extraction

The automation of DNA-PAINT for NPs generates large datasets that need to be analyzed to quantify, for example, the active ligands on the particles’ surface by qPAINT. While counting localization events per particle provides a relative measure of NP functionalization, qPAINT allows for absolute quantification of ligands.^10^ This is not trivial when a large amount of data is produced, which requires an automated batch analysis pipeline. In this section, we describe such an automated analysis pipeline and the resulting NP features extracted from our analysis.

As a proof-of-concept, this was applied to 500 nm polystyrene NPs functionalized with three different DNA docking strands through biotin-streptavidin binding. These were measured with various relative degrees of functionalization, i.e., having different amounts of DNA attached. Prior estimates of functionalization from earlier measurements were used to guide simulation input parameters for optimizing experimental conditions (Supporting Figure 5).

The analysis is largely automated and is divided into two steps. First, pre-processing was performed in THUNDERSTORM^24^ followed by qPAINT analysis and feature extraction using the app nanoFeatures.^25^ The pre-processing, which can be run through a macro if desired, consists of fiducial-based drift correction and minor density filtering. The density filtering gets rid of non-specific background localizations and significantly improves processing speed for the subsequent clustering step.

The resulting files were then processed using the MATLAB GUI nanoFeatures, previously developed in the group. This allows for alignment of the different color channels based on fiducial positions, and then runs a DBScan clustering algorithm, followed by filtering and qPAINT analysis to provide the final list of features as described in the Materials and Methods. The app aligns localizations from all channels, based on fiducial files input by the user. Then it uses the DBScan algorithm to cluster the localizations and applies user-defined filters based on radius and localization count thresholds. The qPAINT analysis generates the timetraces for the individual clusters for each imager, and can also be set to apply a non-specific filter to exclude clusters lacking consistent binding events. Final values are calculated based on the kinetic parameters and imager concentrations that are input, and the true mean dark time obtained by fitting the cumulative distribution function of the dark times for individual clusters.

Figure 4 shows the results of three different NPs analyzed with this pipeline. The obtained mean values match well with expectations based on known functionalization for 1 mol DNA/g polystyrene (Supporting Figure 6). Variability between samples can be explained by a combination of user error, instrument inaccuracy, and statistical variation. Consider, for example, that pipettes used ot prepare the particles often carry an inherent error of 1-2%. Notably, the distributions reveal significant heterogeneity, which is not easily deduced by bulk characterization techniques, but becomes abundantly clear when employing single-molecule techniques like DNA-PAINT. Population variance can be substantial, and it is not equal between every sample either, which is critical in contexts like drug delivery. Differences in the molecular abundance of target molecules can have a significant impact on the eventual functionality. This provides further argument for the necessity of single-molecule characterization techniques to properly characterize nanoparticles.

**Figure 4.**
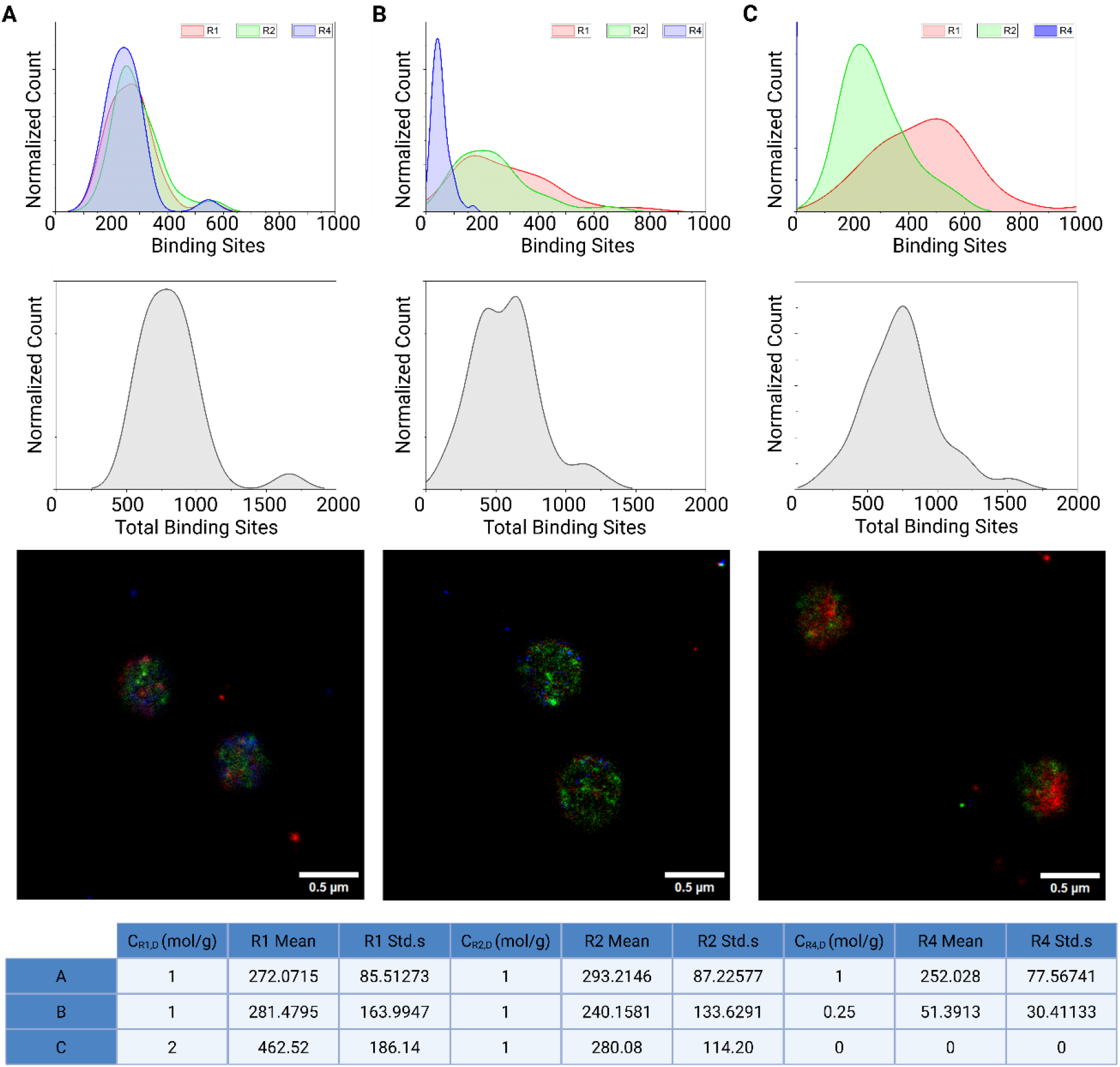
Example results of the multiplexed automated imaging for three different samples (A, B, C). From top to bottom, the figure shows: the distribution of the calculated binding sites for all three imager channels; the distribution of total binding sites per particle; visual examples of reconstructed particles colored per imager; and finally, a table of results showing the docking concentration used and the mean and standard deviation obtained for each imager.

In principle, this methodology can be extended to different particles. To explore the generalizability to soft particles, we applied the workflow to cholesterol-tagged liposomes (Fig. 5). Lipid particles are of particular interest due to their advantages as drug carriers.^26,27^ Here, these particles were mixed with cholesterol-DNA with a TEG linker to incorporate the docking DNA (see Materials and Methods) as a substitute for functional cholesterol-conjugated ligands.

**Figure 5.**
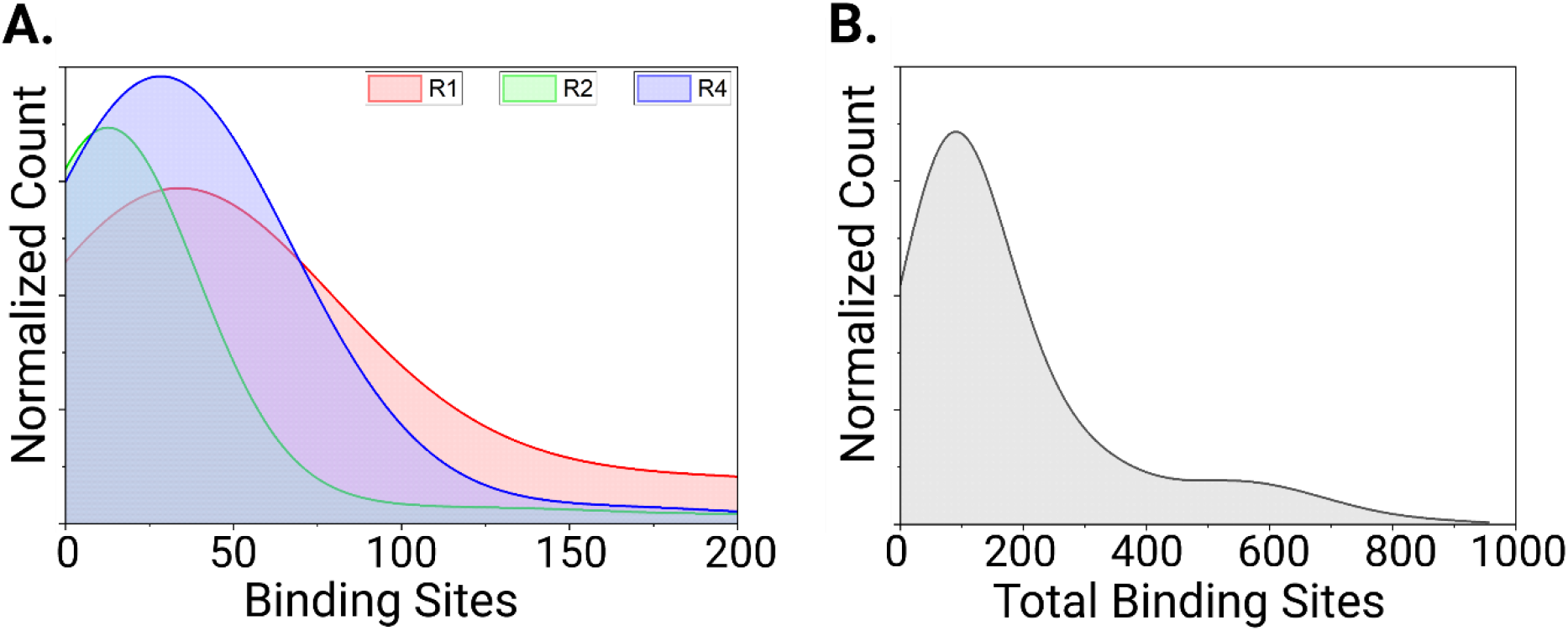
qPAINT result of multiplexed DNA-PAINT on 100nm extruded liposomes (99.5% DOPC, 0.5% DSPE-PEG-Biotin) with three different Cholesterol-TEG-ssDNA docking strands, showing the (A) binding sites per DNA sequence and (B) the total binding sites.

This proved to be much more challenging, however, as incorporation was far lower than expected. Based on the extruded size and ratio of lipids:cholesterol, the expected amount of DNA/liposome was around 800 per docking strand (see Materials and Methods). Measurements showed an order of magnitude lower incorporation than that (Figure 5). Several factors could be the cause of this inefficiency. The TEG linker increases the hydrophilicity of the molecule, which does aid in the solubility, but may also make it less energetically favorable for the insertion into the lipid bilayer. This is particularly relevant as the incorporation was done by passive incubation. The chosen lipid, DOPC, is also highly fluid, which eases sample preparation, but may also not favor stable anchoring of the DNA strands.

This underscores the importance of per-particle quantification, especially in translational nanomedicine, where ligand density often dictates efficacy. Counting capability is essential for estimating functionalization efficiency, which can be dramatically lower than expected based on input reagents, as shown with the liposome case. Equally important is quantification per particle, given that heterogeneity can be quite significant, as demonstrated in all measurements in this work. Only a subset of the population is likely to carry the desired number of functional ligands, and this fraction decreases when multiple ligands are involved. To improve the clinical translation of nanomedicine, single-particle characterization and improved understanding of heterogeneity in particle functionalization are key.

## Conclusion

We have introduced a workflow to enable automated, multiplexed DNA-PAINT experiments for the characterization of nanoparticles at the single-molecule level. This approach combines simulations, imaging, and data analysis into one pipeline, significantly enhancing throughput, allowing for high-resolution imaging that provides detailed insights in a practical timeframe. The methodology is highly flexible, making it compatible with a range of flow and microscope systems. This adaptability means that labs that are already equipped with high-resolution microscopy systems can implement this workflow at minimal cost, thus expanding the accessibility of single-molecule nanoparticle characterization. Future integration with machine learning-based feedback could further automate parameter optimization, bringing high-content single-molecule characterization in line with self-driving lab principles.

We have further demonstrated how critical proper experimental parameters are for the accurate quantification of ligands. Computational simulations play an important role here, as they allow researchers to fine-tune experimental conditions without extensive trial-and-error. Correctly identifying the proper parameters can set one’s experiments up for success, ensuring reproducible and accurate results even for complex samples or low-density samples.

Furthermore, our proof-of-concept highlights the value of qPAINT, particularly the ability to observe heterogeneity within nanoparticle populations that would otherwise be obscured by conventional bulk techniques. Intrasample variation can dramatically impact particle function, and uncovering these differences will be a key factor in identifying effective nanoparticle formulations. The ability of qPAINT to directly quantify the number of ligands on nanoparticles further enhances its utility, offering precise information that can be applied to downstream applications, such as targeted drug delivery or materials science research. This methodology can be readily expanded by coupling docking strands to antibodies, for example, or by applying this to more relevant particle types such as liposomes.

In the future, this work could be extended by incorporating real-time analysis to dynamically adjust acquisition parameters based on previously run simulations, further increasing throughput and reducing data redundancy. The modular nature of the workflow also lends itself to integration with automated sample preparation systems or downstream assays, enabling closed-loop design–test cycles for nanoparticle development. Altogether, the automated workflow and granular insights provided by this platform offer a versatile tool for both fundamental studies and translational applications.

## Materials & Methods

### Materials

DNA strands, functionalized with either Biotin or a TEG-Cholesterol linker, were obtained from IDT and selected based on previous work by Jungmann et al.^23^ (Supporting Table 1). Polystyrene particles with streptavidin of approximately 500-600nm were purchased from Spherotec (SVP-05-10). PLL-PEG with 50% biotin (PLL(20)-g[3.5]-PEG(2)/PEG(3.4)-biotin(50%)) was purchases from SuSoS AG. Phosphate-buffered saline tablets were purchased from Sigma-Aldrich (P4417). Sodium chloride was purchased from Carl Roth GmbH (P029.1). The 18:1 DOPC lipid (1,2-dioleoyl-sn-glycero-3-phosphocholine; 850375C), DSPE-PEG-Biotin(2000) (1,2-distearoyl-sn-glycero-3-phosphoethanolamine-N-[biotinyl(polyethylene glycol)-2000] (ammonium salt); 880129C), 10mm extrusion filter supports (6100014-1EA), PC membranes (0.1um 610005 and 0.2um 610006) and extruder set (610000) were obtained from Avanti Research. Chloroform (1077024) was purchased from Merck. The 1M 4-(2-hydroxyethyl)-1-piperazineethanesulfonic acid (HEPES) buffer (15630080) was obtained from Thermo Fisher Scientific.

Imaging buffer was prepared using 1xPBS, 500mM NaCl, and 1mM EDTA at pH 7.4, to stay consistent with kinetics reported by Jungmann.

### Timetrace simulations

Time versus intensity traces of a certain duration (T) are simulated for a single particle in a region of 8 by 8 pixels. All intensity values shown are pixel averages, rather than the total intensity of the area. The particle has a specified number of binding sites (Nb), which is considered to be the ground truth. The mean dark and bright times are calculated from the imager concentration plus either the second-order association constant (k_on_) or the first-order dissociation constant (k_off_). As has been previously shown, bright and dark times are exponentially distributed^17^. Therefore, the calculated mean dark and bright times are used to generate a randomized exponential distribution, which can then be used to generate the timetraces. Matlab has an in-built command (*exprnd)*, which creates a list of exponentially distributed random dark and bright times based on the calculated means. These values are concatenated, but the values are specified down to 1/10^th^ of a millisecond. To ensure proper sampling, the model takes a sampling frequency equivalent to 1/100^th^ of the shortest of either the dark or bright times. For each time step, it is assigned an intensity value depending on whether it falls within a dark or a bright time.

Dark times are assigned an initial intensity value of 0, while bright times are assigned an intensity value dependent on the laser power (P). Based on experimental data, we know that intensity is not consistent between events. Photon adsorption and emission rates follow Poisson statistics, and neither particles nor sample surfaces are fully homogeneous. This usually leads to a log-normal distribution, as shown by Kish et al^28^. The in-built MATLAB function *lognrnd* is used to generate a list of random intensities in a log-normal distribution. The mean intensity scales with the laser power (P) and was based on experimental data obtained at a camera frequency of 10 Hz. When binning the time trace to the set camera frequency (F), this is taken into account and corrected for, so that we properly account for events that start within the duration of a frame.

Lastly, the background noise due to the free imager and camera base level is taken into account. The camera used here has a baseline of 400, and the background intensity level is known to scale with imager concentration, camera frequency, and laser power based on previous measurements. The MATLAB function *normrnd* is used to generate a Gaussian distribution of background intensities, which is combined with the event timetraces to construct the total time trace. If parameters like the laser power or imager strand concentration are set extremely high, pixels may become saturated, and they are capped out at 2^16^.

### Accuracy and precision calculations

Timetrace simulations are run for a combination of different measurement times (T) and imager concentrations (ci), while keeping all other parameters constant. Each condition is repeated 1000 times to ensure proper statistics. From each timetrace, the dark times are extracted by determining whether they exceed the threshold, set as 4 times the standard deviation of the background. Single-frame dark times are removed, as this is also done in our actual later analysis. This is because single off-frames are more often caused by flickering, which means fluorophores are sometimes not detected for a brief moment, leading to inaccuracy. A cumulative distribution function of the dark times is generated and then fitted to obtain the true mean dark time (τ_d_), which is used to calculate the number of binding sites (N_b,out_). Then, these values are used to obtain the precision defined as (1-(std(N_b,out_)/mean(N_b,out_), and the accuracy defined as (1-|(N_b_-N_b,out_)/N_b_) for each condition. They are defined such that values closer to 1 are more accurate and precise. The results are compiled in a matrix, which is then plotted as two heatmaps. The heatmaps are centered around 90% and scaled such that it is easy to identify that specific cut-off. For interpretation, we consider results that have both an accuracy% and precision% over 90% to be sufficient for determining our experimental parameters.

### Sample preparation

Functionalized polystyrene particles are prepared by mixing 5 μL of 10 mg/mL streptavidin-coated polystyrene particles with a mixture of biotin-docking strands at 10 μM, with the amount of DNA varying per experiment. The sample is then left to stir in a Thermoshaker at RT for 1 hour. It is diluted to 200 μL total volume with PBS (0.25 mg/mL particle). To remove free DNA, it is subsequently spun down at 16k RPM at 4C for 10 minutes. The solvent is taken off, and the particles are then resuspended. This is repeated 3 times. The particles can then be stored at 4 °C in the fridge for several weeks.

Samples are prepared in an Ibidi VI 0.5 μ-Slide with glass bottom (Ibidi GmbH, 80607). The slide is ozone cleaned before surface functionalization in a UV Ozone Cleaner (Novascan PSD series, Digital UV Ozone System) for 15 minutes. First, the channel is filled with 40 μL of diluted 100nm Tetraspeck beads (ThermoFisher Scientific Inc., T7279) and left to incubate for 15 minutes. The channel is then washed with 70 μL of PBS and filled with 40 μL of 0.1 mg/mL PLL-PEG-biotin and incubated for 15 minutes. The channel is then washed with 70 μL of PBS and filled with 40 μL of the particle sample at 0.25 mg/mL, then left to incubate for 30 minutes. Finally, it is washed once more with 70 μL of PBS and then partially filled with PBS to prevent air bubbles in later flow steps that would otherwise form due to the air volume in the reservoir.

Liposome particles were prepared by manual extrusion. Mixtures of 10mM DOPC and DSPE-PEG-Biotin were prepared in advance by dissolving them in chloroform. A glass vial was cleaned with acetone and dried with nitrogen. Then 199 μL of 10 mM DOPC and 1 μL of 10 mM DSPE-PEG-Biotin were added to end up with a mixture of 99.5% DOPC and 0.5% DSPE-PEG-Biotin. This was vortexed while evaporating with nitrogen to form a thin lipid film. It was then redissolved in 1 mL buffer consisting of 1mM HEPES and 50 mM NaCl. It was subsequently extruded 21 times by pushing it through 0.2 μm PC membranes, using an extrusion block and two syringes. This was then followed by an additional extrusion step of 21 repetitions through 0.1 μm PC membranes. The remaining sample is transferred to an Eppendorf and then stored in the fridge and used within 24 hours.

To determine the amount of DNA per liposome, we need to know the number of lipid molecules in a DOPC liposome. Assuming that they form a single lipid bilayer with head groups on each side, we can simply calculate the total internal and external surface area and divide by the lipid head group area. In this case, the amount of DOPC lipid per liposome is 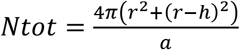, where r is the liposome radius, h is the bilayer thickness (assumed to be 5 nm), and a is the lipid head group area (0.71nm for DOPC). For 100nm DOPC liposomes that we extrude to, this gives 78976 head groups. Thus, to obtain 800 DNA/liposome, we require a roughly 1:100 ratio, and add 10 μL of 100uM of each cholesterol-DNA to 50uL of 2mM DOPC liposomes. This is left to incubate on a shaker for 1.5 hours, and then diluted 1000 times for the final measurement to obtain a reasonable particle density.

Sample preparation for the liposome experiments is the same as above, except after the PLL-PEG-biotin incubation and washing step, there is an additional incubation with 0.1 mg/mL streptavidin for 15 minutes. The slide is then washed, and 40 μL of the diluted liposome sample is added and left to incubate for 30 minutes. Finally, it is washed once more with 70 μL of PBS and then partially filled with PBS to prevent air bubbles in later flow steps that would otherwise form due to the air volume in the reservoir.

### Flow set-up

For the flow set-up, the LSPOne Pump by Advanced Microfluidics was used with a 1 mL syringe and 10 valve ports (AF-LSP-D10-05-UK). The slide was connected to the pump using Elbow Luer Connectors (Ibidi GmBH, 10802), and 1/16” OD x 0.5 mm ID PTFE tubing (BL-PTFE-1602-20) and flat-bottom fittings (CIL XP-245X) by Darwin Microfluidics were used to connect to the sample as well as all the solutions, which were kept in either falcon tubes or DNA LoBind eppendorfs. All tubing was measured to obtain the internal volume. Before starting a measurement, the tubing connecting to the buffer and the imager solutions is first primed by drawing up slightly more than their internal volume and then dispensing to the waste. Then buffer is dispensed to the tubing, which will be later connected to the sample, until it is filled and a slight droplet forms at the edge of the luer connector, before connecting with the ibidi slide. This is to avoid a large air bubble passing through during the first flow step, which would potentially devastate the sample.

The LSPOne Pump can be controlled using ASCII-based text commands as listed in the official documentation. During the measurement, the pump is controlled by a custom MATLAB GUI, which can construct and send these commands to achieve the desired action. It can manually move between the different valve ports and pick up and dispense specific volumes at different speeds. More importantly, it incorporates a protocol reader that can read in text files to control the pump, making it trivial to write expansive experiment workflows. It can also set up actions to happen at specific timepoints, counting from the start of the protocol. This allows us to set up flow steps to happen in between imaging steps, without user interference.

Generally, the sample volume is chosen to ensure that the volume in the channel is fully displaced, to avoid dilution of the imager. All the tubing was measured, and based on the internal diameter, we know that the volume of the output line is roughly 240 μL, and the volume for the line going from the sample to the waste is 180 μL. The ibidi slide has a channel volume of 40 μL, but each of the reservoirs has an additional volume of 60 μL. This gives a total volume from output to waste of 580 μL. We flow a volume of 800 μL across to make sure that it is fully displaced.

In terms of flow speed, we take up and dispense volumes at 30 uL/s, which keeps the flow steps short without generating too much pressure, which would pop off the luer connectors. We observe that all particles remain at these flow speeds, and slightly higher ones even.

### Imaging set-up

Images were recorded in an Oxford Nanoimager microscope (ONI, Oxford, UK). The imaging was carried out in total internal fluorescence (TIRF) configuration with a 100x 1.4 NA oil-immersion objective. Imagers were acquired on a 428 by 428 pixel area, with a pixel size of 0.117 μm on a sCMOS camera. All images were recorded at 50 ms exposure time for 500 seconds, and illuminated with a 640nm laser at 30mW output. The in-built Python console was used to set up an automated imaging workflow consisting of repeated multiacquisitions that recorded three fields of view per imager, with a waiting step in between multiacquisitions to account for the flow steps. Schematically, an example workflow looks as follows:

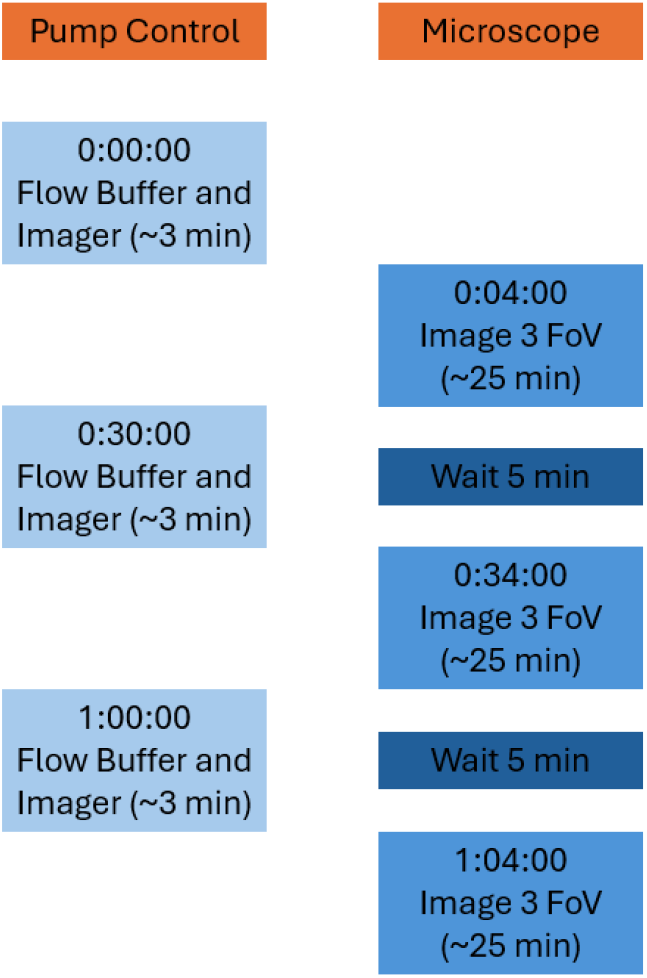

A certain amount of wiggle room is kept for every step to avoid overlap in case either instrument experiences a few seconds delay for whatever reason.

### Analysis workflow

The Nanoimager software has an in-built function that was used to do the Gaussian fitting and obtain the localizations, before outputting them in a CSV. If other instrumentation is used, this can easily be replaced by running the analysis of a TIFF or ND2 file in THUNDERSTORM.^24^ The output was run through a MATLAB script to change the headers to make them compatible with THUNDERSTORM, which was used to run the drift correction and the density filtering. Two files were exported from THUNDERSTORM. One CSV with all the data with slight density filtering to remove background, and one with an extreme filter of 9000 localizations within a 50nm radius that should only retain clusters that are near permanently on, i.e., the fiducials. These fiducial files are important for later alignment.

The rest of the analysis is run in nanoFeatures,^25^ which aligns the images based on the fiducial coordinates, and then clusters and filters particles based on a set of input parameters, including size filtering, a minimum number of localizations, and a non-specific filter, among others. It also runs the qPAINT analysis to obtain the bright and dark times and number of binding sites for each imager channel per particle.

Output is given in a .csv which contains the following features per particle: diameter (nm), aspect ratio, x center coordinate, y center coordinate, cluster localizations; and then for each channel per particle: channel localizations, true mean dark time, R^2^ of fit, mean dark time, median dark time, standard deviation of dark times, mean bright time, median bright time, standard deviation of bright times and calculated target count.

## Acknowledgements

The authors thank A. Daeniker for support with the LSPOne Interface. The authors also thank M. Pontier for initial work on the DNA-PAINT simulations. Schematic illustrations were created using Biorender.com. This work has received funding from the European Union’s Horizon 2020 Research and Innovation program under Grant agreement No 964386—FET Open RIA project acronym “MimicKEY”.

## Code Availability

The code used in this manuscript to perform the simulations was written in MATLAB (v.2020b); the GUI to control the pump was also written in MATLAB (v.2020b); and the scripts to control the microscope were written in Python. All of these are made publicly available at https://github.com/smavanveen/HT-Multiplexed-PAINT.

