## Supporting Information for "Automated and Simulation-Guided Multiplexed DNA-PAINT for Nanoparticle Characterization"

| ID | Docking Sequence | Imager Sequence | $k_{on}$ | $k_{off}$ |
|---|---|---|---|---|
| **R1** | 5'-/5Biosg/TCC TCC TTT-3'<br>or<br>5'-/5CholTEG/TCC TCC TTT-3' | 5'-AGG AGG A/3ATTO633N/-3' | 7.8*10^6 | 4.7619 |
| **R2** | 5'-/5Biosg/ACC ACC AAA-3'<br>or<br>5'-/5CholTEG/ACC ACC AAA-3' | 5'-TGG TGG T/3Atto633N/-3' | 9.3*10^6 | 1.6129 |
| **R4** | 5'-/5Biosg/ACA CAC AAC-3'<br>or<br>5'-/5CholTEG/ACA CAC AAC-3' | 5'TGT GTG T/3Atto633N/-3' | 5.5*10^6 | 4.1666 |

*Supporting Table 1 Sequences and kinetic data used in experimental design, based on those reported by Jungmann et al..*[23]

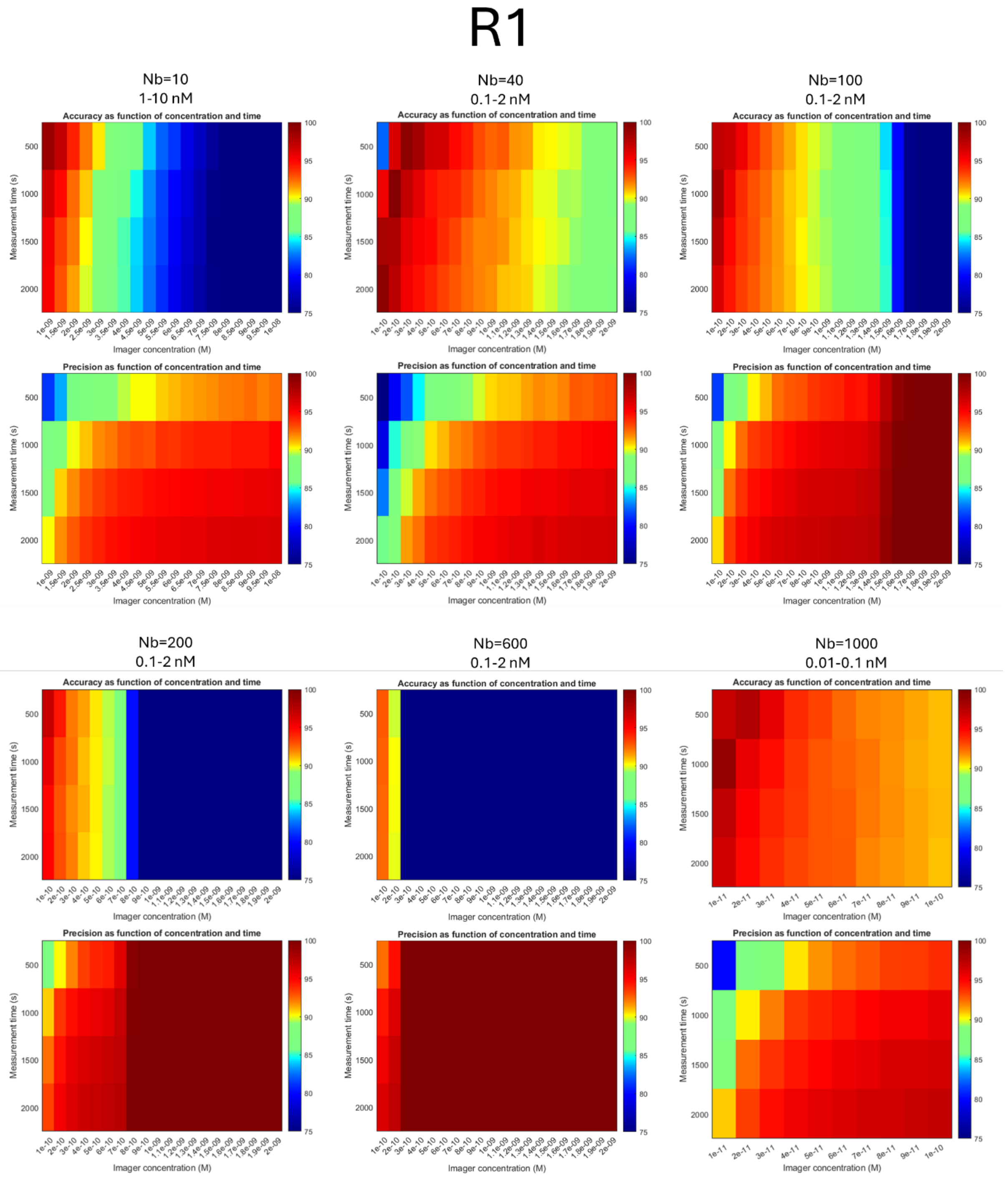


*Supporting Figure 1 Heatmaps of accuracy and precision for R1 imager-docking pair at various concentrations, for different amounts of binding sites (Nb). Each combination was ran for 1000 repetitions.*

# R2

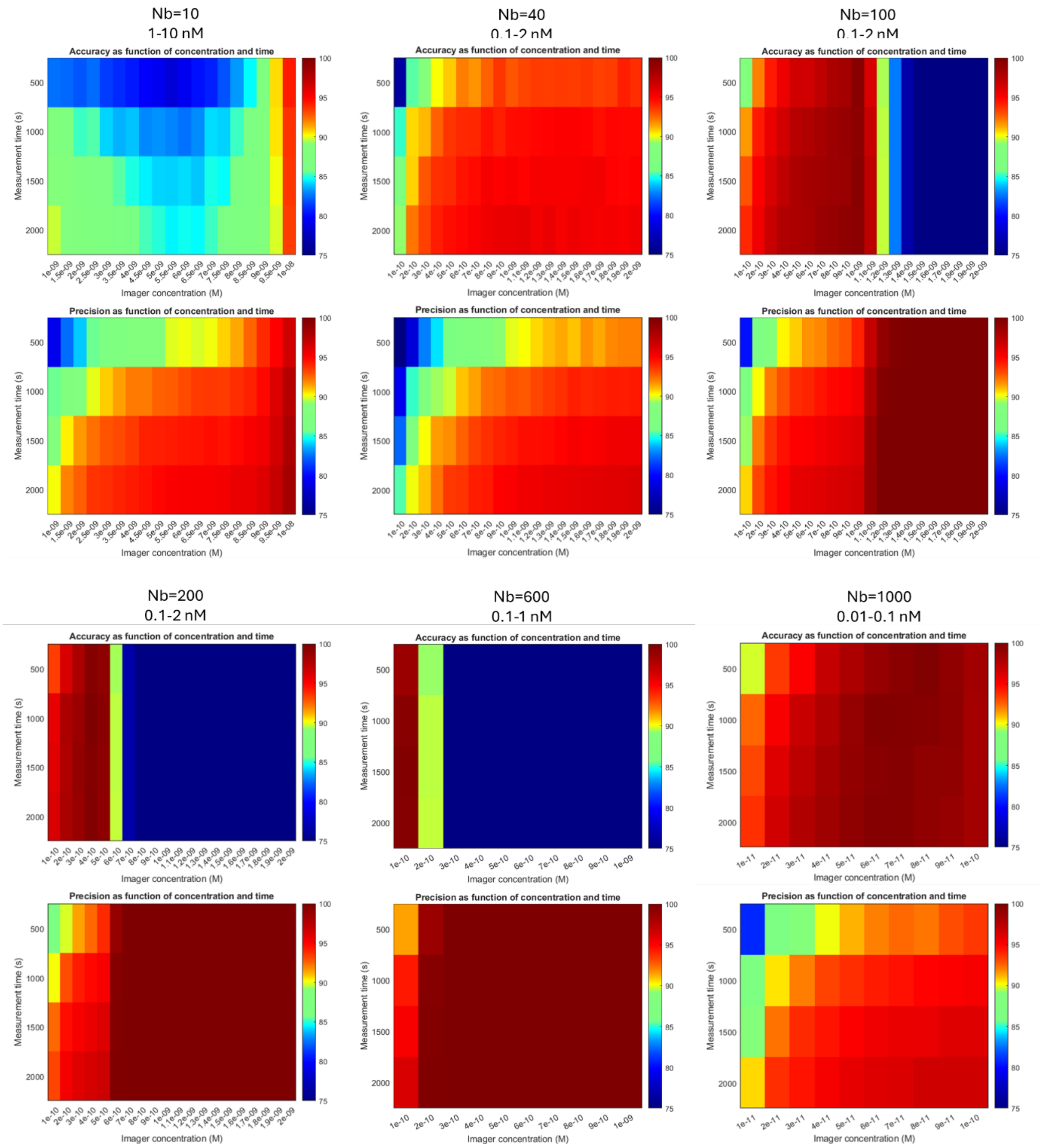


*Supporting Figure 2 Heatmaps of accuracy and precision for R2 imager-docking pair at various concentrations, for different amounts of binding sites (Nb). Each combination was ran for 1000 repetitions.*

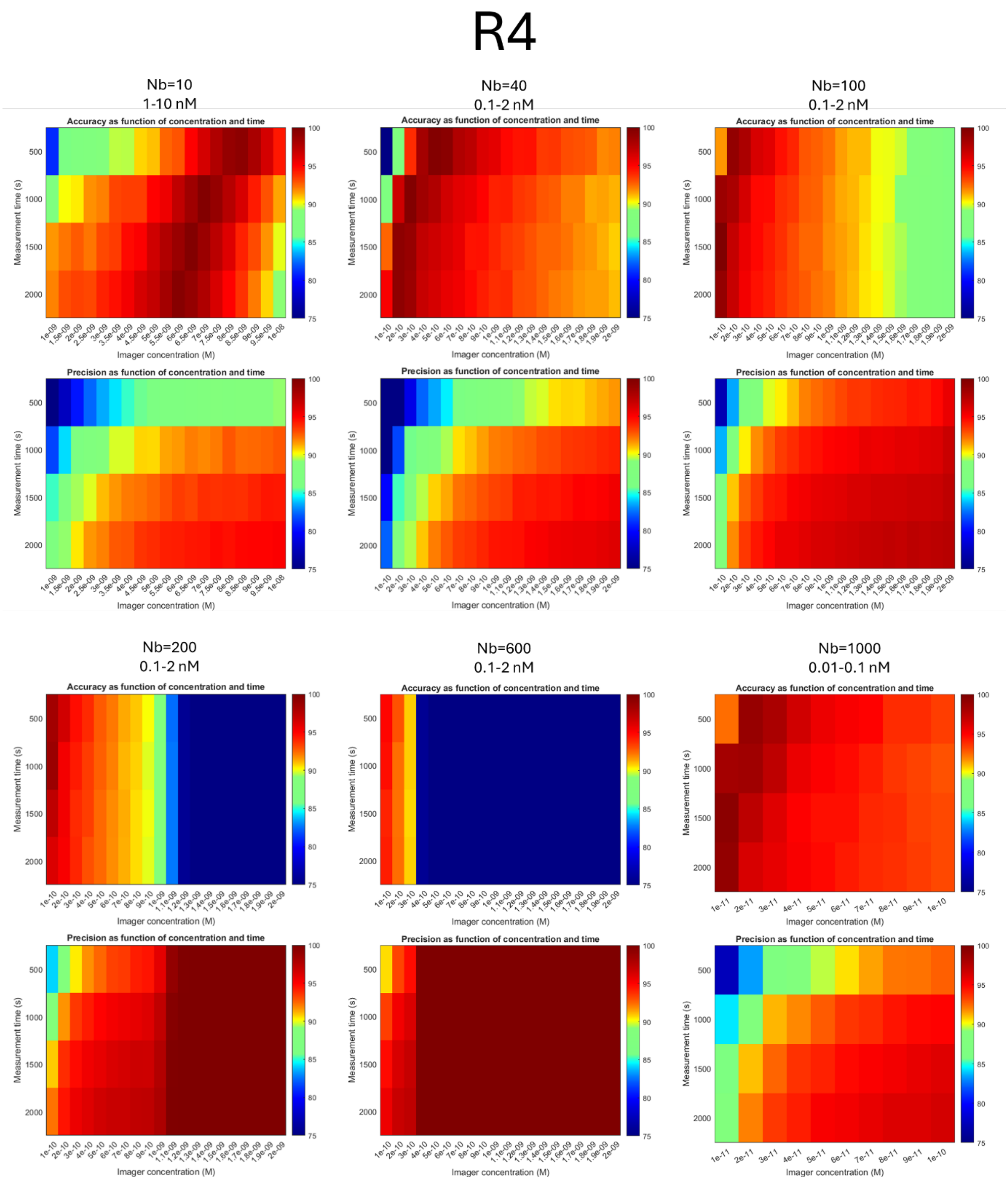


*Supporting Figure 3 Heatmaps of accuracy and precision for the R4 imager-docking pair at various concentrations, for different amounts of binding sites (Nb). Each combination was ran for 1000 repetitions.*

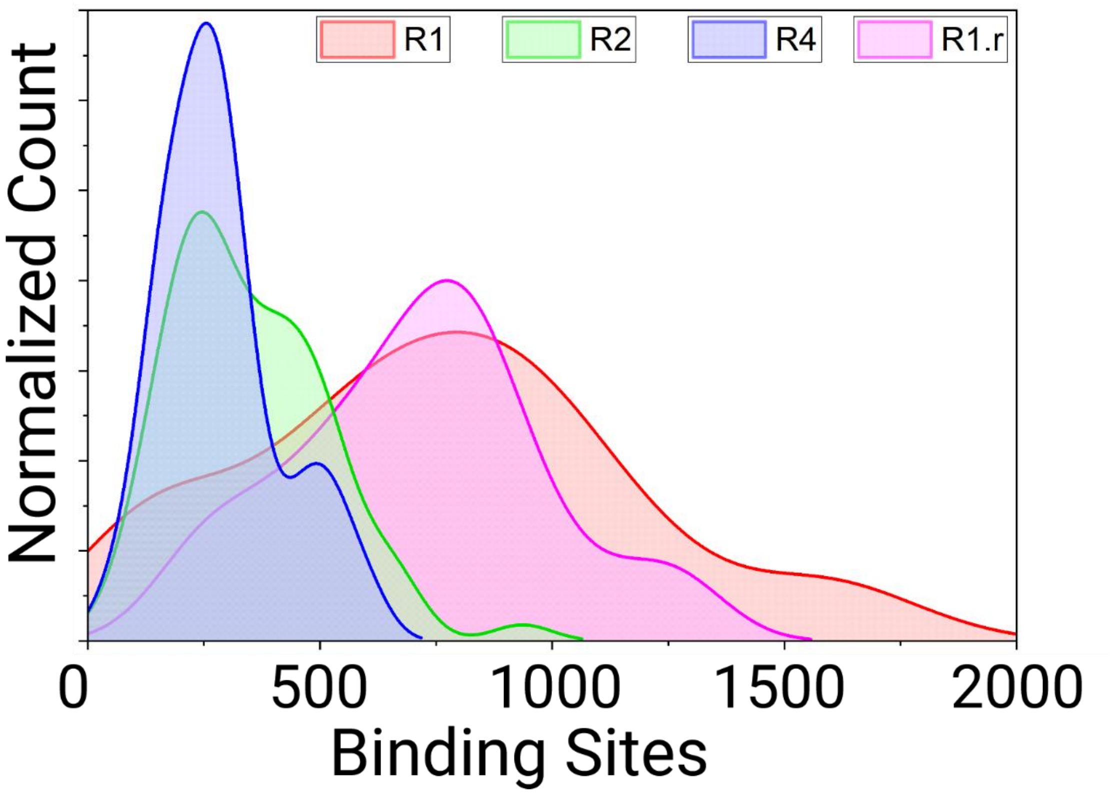


*Supporting Figure 4 Binding sites as measured by qPAINT for the measurement referenced in Figure 3, with R1 imager repeated at the end as a control. Ratios of docking added were 2:1:1 for R1:R2:R4. Obtained means are (R1=759, R2=348, R4=285, R1_repeat=716). The slight difference with the repeat falls within what can reasonably be differences in accuracy of counting at this timescale.*

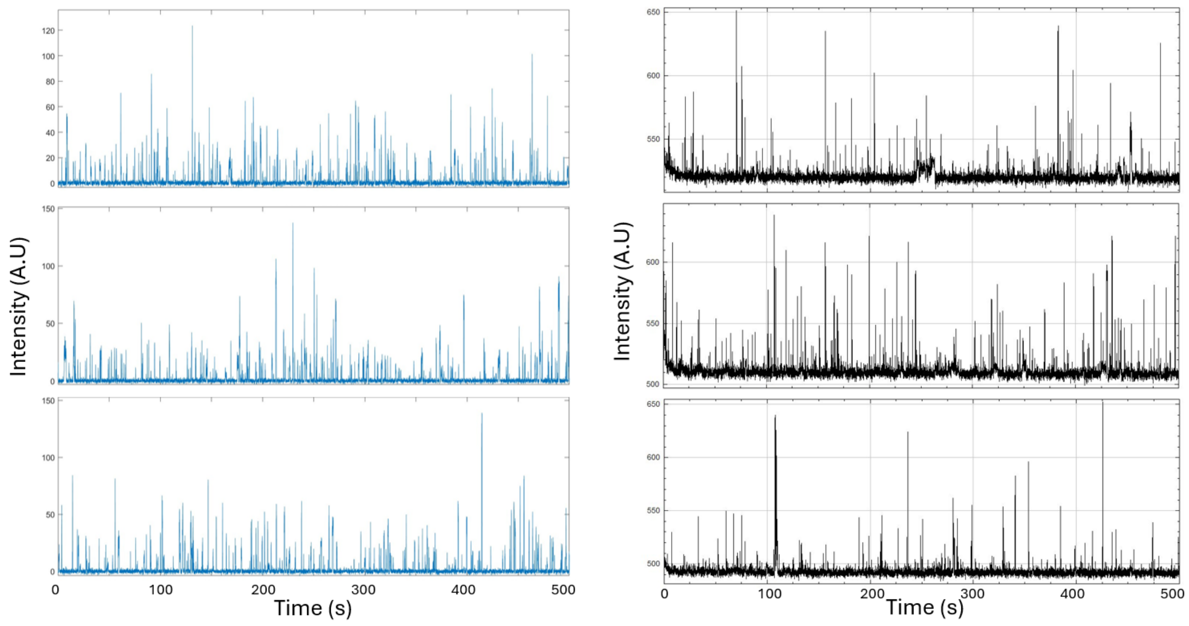


*Supporting Figure 5 Comparison of simulated timetraces (left) versus the timetraces observed in a real measurement for the same parameters (right). The timetraces were determined for an 8x8 pixel area, for 0.5 nM R4 imager strand on particles with (on average) 250 binding sites.*

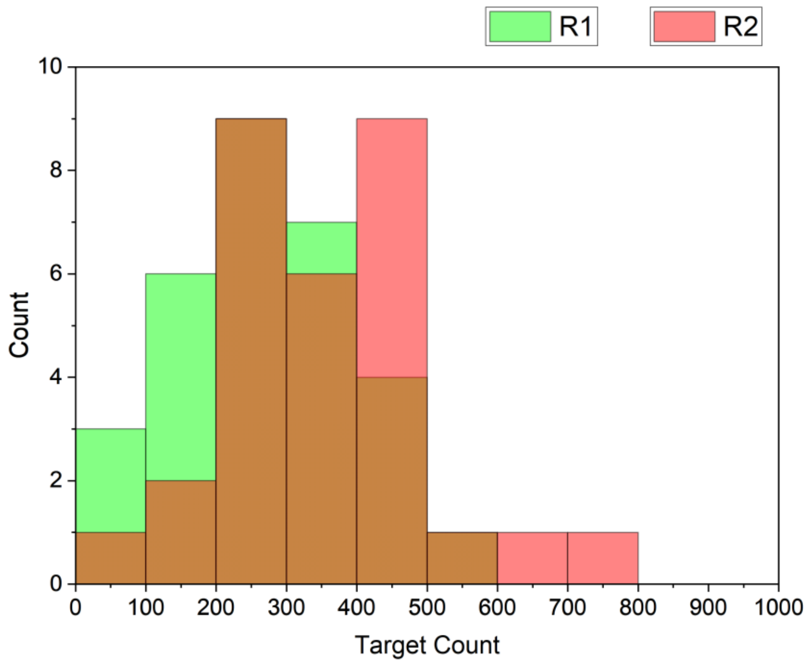


*Supporting Figure 6 Quantitative result of one of the initial dual-color experiments run on polystyrene-DNA particles. For docking concentrations of 1mol/g, obtained means were (R1=270, R2=330).*
